# Protein Stability, Turnover Kinetics, and Abundance Constrain the Scaling of Protein Interaction Networks

**DOI:** 10.64898/2026.05.11.724303

**Authors:** Muskan Goel, Daniel A. Nissley, Xavier Castellanos-Girouard, Charles P. Kuntz, Yiqing Wang, M. Shahid Mukhtar, Adrian Serohijos, Jonathan P. Schlebach

**Author notes:** Authors contributed equally (Muskan Goel, Daniel A. Nissley).

## Abstract

The propensity of proteins to form oligomers is ultimately dictated by their structural configuration(s). Proteins that persist in a discrete conformational state may form a limited number of specific interactions while those that sample a broader structural ensemble may instead associate with a wider array of partners. These intrinsic tendencies potentially constrain the way proteins navigate wider interaction networks. In this work, we aggregated and surveyed a wide variety of biophysical, biochemical, and cellular descriptors of the *S. cerevisiae* proteome to identify biases in the connectivity of its protein-protein interaction network. Using mass spectrometry-based interactome measurements and various protein stability estimates, we find that a disproportionate number of abundant, yet unstable binding proteins act as network hubs. Moreover, we show that these features alone can be used to discriminate between hubs and non-hub proteins with high accuracy (AUROC = 0.898). Interestingly, we find that half-lives of hub proteins depend on whether or not they reside within static complexes and/ or whether they interact with molecular chaperones. Finally, we note that the observed connectivity biases associated with abundant, unstable proteins only pertain to network hubs, but not to the bottlenecks that connect them. Together, our findings reveal how the conformational stability of a protein may constrain its context within protein-protein interaction networks.

## Introduction

Protein-protein interactions (PPIs) emerge from the intrinsic biophysical properties of the proteins and the physicochemical context of their environment. To wit- the conformational stability and dynamics of proteins ultimately govern their competence for binding,^1^ while protein abundance sets the energetic balance between monomers and oligomers.^2^ These biophysical principles provide a deterministic lens to rationalize the factors that drive one protein to interact with another. Computational models have provided suggestions as to how genetic modifications shape physical interaction networks, and vice versa.^3–7^ However, efforts to survey how folding and binding energetics influence PPIs in the cell, where proteins can interact with thousands of different potential binding partners, remains fundamentally challenging. Nevertheless, new scalable tools to rationalize the biophysical properties of proteins combined with mounting experimental descriptors of PPI networks are providing new opportunities to explore their physical constraints.

At the cellular scale, the emergent scaling of PPI networks appears to endow them with collective robustness to genetic and environmental insults.^8–10^ Efforts to map these networks with different experimental tools have provided a mix of convergent and divergent perspectives on their finer topological properties. Despite poor overlap between their specific edges, network models reconstructed from binary yeast two-hybrid (Y2H) measurements and proteomic affinity enrichment mass spectrometry (AE-MS) exhibit heavy-tailed degree distributions that can be approximated by power laws.^11–15^ Efforts to explore these networks using distinct methodologies have revealed several principles in biological organization.^16–22^ This emerging network view of PPIs revealed that most proteins have only a few interaction partners, with relatively few highly connected proteins serving as network hubs. The connections between these hubs, which are mediated by bottleneck proteins, connect functional modules that support core cellular processes within the cell.^23^

The collective activity of PPI networks is a selected trait that drives cellular fitness.^24–26^ Nevertheless, it is difficult to understand how evolutionary pressures act on this network without a full understanding of the means by which the collective energetics of protein ensembles give rise to cellular interactomes. While scalable approaches to probe PPI networks have been pioneered over the past few decades, efforts to probe the biophysical properties of the interactors has historically required focused investigations of individual proteins. However, advances in high throughput biophysics^27^ and structural modeling^28–31^ in the past decade are beginning to open new doors for the investigation of protein biophysics at scale. In conjunction with the emergence of highly comprehensive interactomes,^15^ proteome-wide surveys of protein abundance,^32,33^ turnover kinetics, ^34,35^ and thermal stability ^36^ have reached a critical mass that can be paired with biophysical tools to search for the mechanistic underpinnings of proteomic behavior.

In this work, we survey the physical constraints on the *S. cerevisiae* interactome by analyzing its network topology in relation to an array of computational and bioanalytical descriptors of protein abundance, protein turnover, and protein stability measurements. Statistical analyses reveal that network hubs exhibit distinct patterns of abundance, turnover, and stability that can be used to accurately predict which proteins are highly connected within the PPI network. We further find that hub proteins embedded in stable protein complexes (designated herein as static hubs)^37^ typically have higher folding stability relative to those that form many interactions outside of a complex (designated dynamic hubs). Finally, we show that dynamic hub proteins tend to form more interactions with molecular chaperones in a manner that coincides with increased kinetic stability in the cell. We discuss how these relationships help reconcile protein-level constraints with emergent network organization and outline implications for the co-evolution of proteome biophysics and network topology.

## Results

### Network Context of Abundant Proteins

To survey the distribution of biophysical and biochemical properties of proteins across PPI networks, we first used NetworkX to generate two independent models of the *S. cerevisiae* interactome based on either immunoprecipitation mass spectrometry (AE-MS)-based interaction measurements or yeast two-hybrid (Y2H) interaction measurements (Fig. 1A-B). To analyze the network context of each protein, we also calculated a series of eight graphical descriptors for each node that reflect their centrality and/ or betweenness within the network model (Fig. 1C). The available measurements cover over ∼65% of the proteome in the AE-MS network (3,927 nodes), with slightly lower coverage (∼33%) in the Y2H network (2,018 nodes). There were also fewer interactions in the Y2H network (2,930 edges) relative to the AE-MS network (31,004 edges). However, both networks exhibit degree distributions that are reasonably well approximated by a power law (Fig. 1D). These observations suggest both network models have a relatively low proportion of high-degree nodes (i.e. hub proteins). Nevertheless, a structural comparison of the subnetworks within these models suggests they share few topological features-only 2.3% of the hubs exhibit similar connectivity (Jaccard similarity index ≥ 0.3, Fig. 1E). This analysis of current interactome data echoes previous investigations highlighting the divergence of Y2H data relative to other interaction measurements.^11–14^

**Figure 1.**
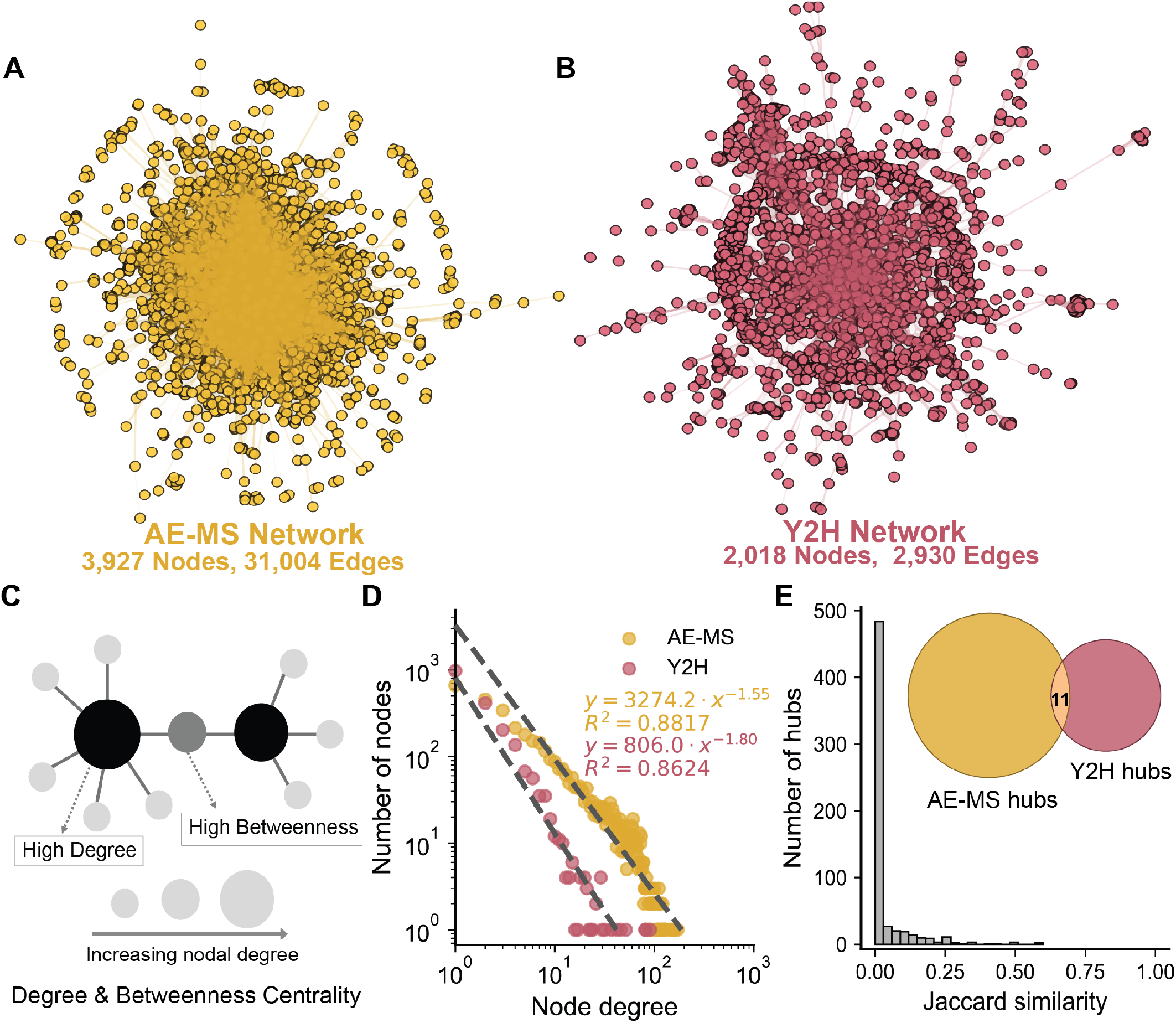
Construction and Characterization of Network Models for the *S. cerevisiae* Interactome. The scaling and similarity of network models derived from collections of proteomic mass spectrometry (AE-MS, yellow) or yeast-two-hybrid (Y2H, pink) interaction measurements are quantitatively compared. Network diagrams depict the interactions between nodes within network models generated using either A) AE-MS data or B) Y2H data. Circular nodes represent proteins and the edges connecting them represent interactions. C) A schematic depicts two classes of centrality descriptors for nodes within the network. High degree nodes act as hubs by forming numerous interactions with other nodes while those with high betweenness act as bottlenecks between hubs. D) The number of nodes is plotted against their corresponding degree centrality. Data from the AE-MS (yellow) and Y2H (pink) networks are fit to a power law function. The fitted functions and their corresponding *R*^2^ values are shown, for reference. E) A histogram depicts the distribution of Jaccard similarity index values for hubs within two networks. Most hubs exhibit very low similarity (close to 0), indicating limited overlap in the identity of high-degree nodes. A Venn diagram illustrates the intersection of hubs defined using a Jaccard similarity cutoff of ≥ 0.3. Only 11 hubs are shared between the two networks at this threshold.

Divergence between AE-MS and Y2H interaction measurements could potentially arise from variations in the abundance of native proteins. Whereas Y2H detects interactions between over-expressed proteins that are colocalized within the nucleus, AE-MS are carried out in the context of complex proteomic mixtures of endogenously expressed proteins. To determine whether variations in natural protein abundance are likely to factor into degree distributions, we compared the relative abundance of hub proteins (i.e ≥ 90^th^ percentile degree centrality) to that of the remaining non-hub proteins. On average, the cellular abundance of AE-MS hubs is 163.45% greater than that of non-hub proteins (Mann-Whitney U-test, *p* = 5.6 × 10^−65^), though the same is not true for Y2H hubs (Fig. 2A). Proteins that form bottlenecks (i.e. nodes with high betweenness centrality) within the AE-MS network are also generally more abundant as well (Mann-Whitney U-test, *p* = 1.9 × 10^−30^), though the same is not true for Y2H bottlenecks (Fig. 2D). Cross-referencing these proteins against cellular turnover measurements also reveals that AE-MS hubs exhibit lower kinetic stability relative to non-hub proteins (Mann-Whitney U-test, *p* = 1.4 × 10^−36^), though this trend is once again not observed among Y2H hubs (Fig. 2B). Bottleneck proteins exhibit no difference in turnover kinetics in either network (Fig. 2E). Overall, the divergent properties of hubs and bottlenecks are most apparent in the context of the AE-MS network, which may arise from the preservation of native stoichiometries in AE-MS experiments (see *Discussion*). Nevertheless, both the hubs and bottlenecks within the AE-MS and Y2H networks are statistically enriched with essential proteins (Fig. 2 C & F). Together, these findings suggest hubs are often essential proteins that tend to be highly abundant and rapidly turned over in the cell.

**Figure 2.**
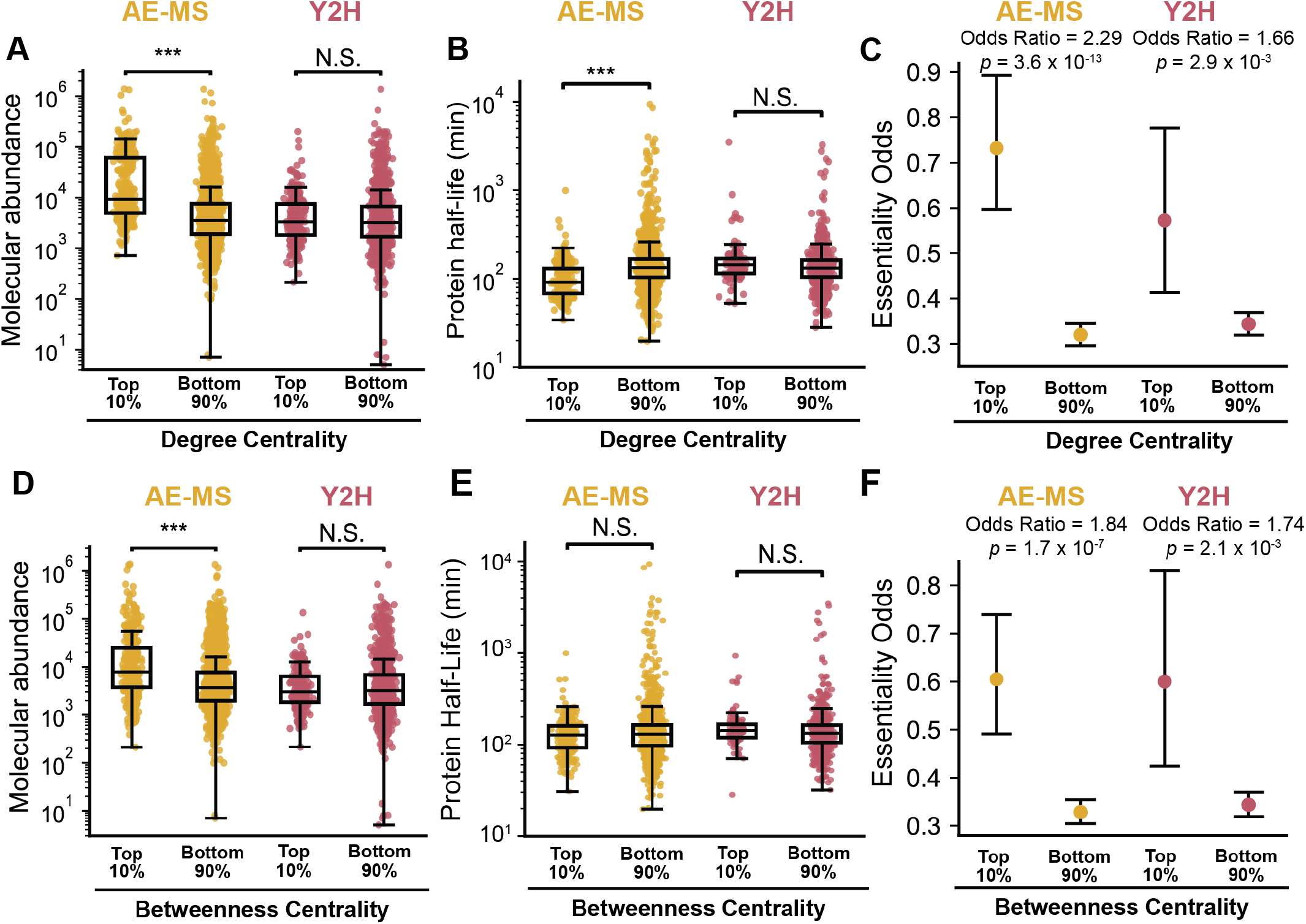
Cellular Abundance and Turnover of Network Hubs and Bottlenecks. The cellular abundance, protein half-life, and essentiality of hub proteins (≥ 90^th^ percentile degree centrality) and bottleneck proteins (≥ 90^th^ percentile betweenness centrality) within the AE-MS (gold) and Y2H (pink) networks are compared. Box and whisker plots depict the statistical distribution of molecular abundance values among the A) hub proteins and D) bottleneck proteins from each network. Box and whisker plots depict the statistical distribution of protein half-life measurements among B) hub proteins and E) bottleneck proteins from each network. The upper and lower edges of the boxes correspond to the 75^th^ and 25^th^ percentile value, respectively. The upper and lower whiskers extend to 1.5 times the interquartile range. Midlines reflect the median value. Statistical significance is indicated by *** (Mann-Whitney U-test, *p* < 10^−30^). The odds that a C) hub protein or F) bottleneck protein is derived from an essential gene is compared to the odds that a non-hub or non-bottleneck within each network, respectively, is essential. Whiskers represent 95% confidence intervals and *p*-values were calculated using Fisher’s Exact Test.

### Network Context of Stable Proteins

Many hub proteins feature an abundance of disordered regions that participate in an array of competing PPIs.^38,39^ These considerations potentially suggest hub proteins may exhibit minimal thermodynamic tendency to adopt a single, ordered conformation. To determine whether hub proteins exhibit bias with respect to protein stability, we generated a variety of protein stability predictions for the yeast proteome and gathered available proteomic stability measurements. We first used a collection of AlphaFold2 models of the yeast proteome in conjunction with a recently described protein stability predictor to generate estimates of the free energy of folding (Δ*G*_f_) for each member of the yeast proteome. We also used these structures to generate stability estimates using both the Rosetta energy function and a previously described length-based scaling function. As these predictions are irrelevant for intrinsically disordered proteins (IDPs) that lack a well-defined native structure, we removed all stability estimates based on AlphaFold2 models that feature a high fraction of residues with low confidence predictions (average pLDDT ≥ 70), which appears to be a strong empirical indicator for intrinsic disorder (Fig. S1). Given that these predictions are also unlikely to be accurate for integral membrane proteins, we used DeepTMHMM^40^ to remove any predictions for proteins that are predicted to contain at least one transmembrane domain. To enhance the accuracy of stability predictions for secreted proteins, we leveraged SignalP predictions^41^ to identify the positions of any signal peptides, then removed these segments from the predicted structures prior to generating refined stability predictions based on structural models lacking the signal peptide (see *Methods*). Finally, to evaluate experimental stability metrics, we also incorporated analyses of previously reported proteomic “meltome” measurements that track the solubility of each protein as a function of temperature. Overall, these stability metrics exhibit considerable variation but generally agree that only a modest fraction of soluble proteins that tend to adopt a discrete native structure lack conformational stability (Fig. 3A-D).

**Figure 3.**
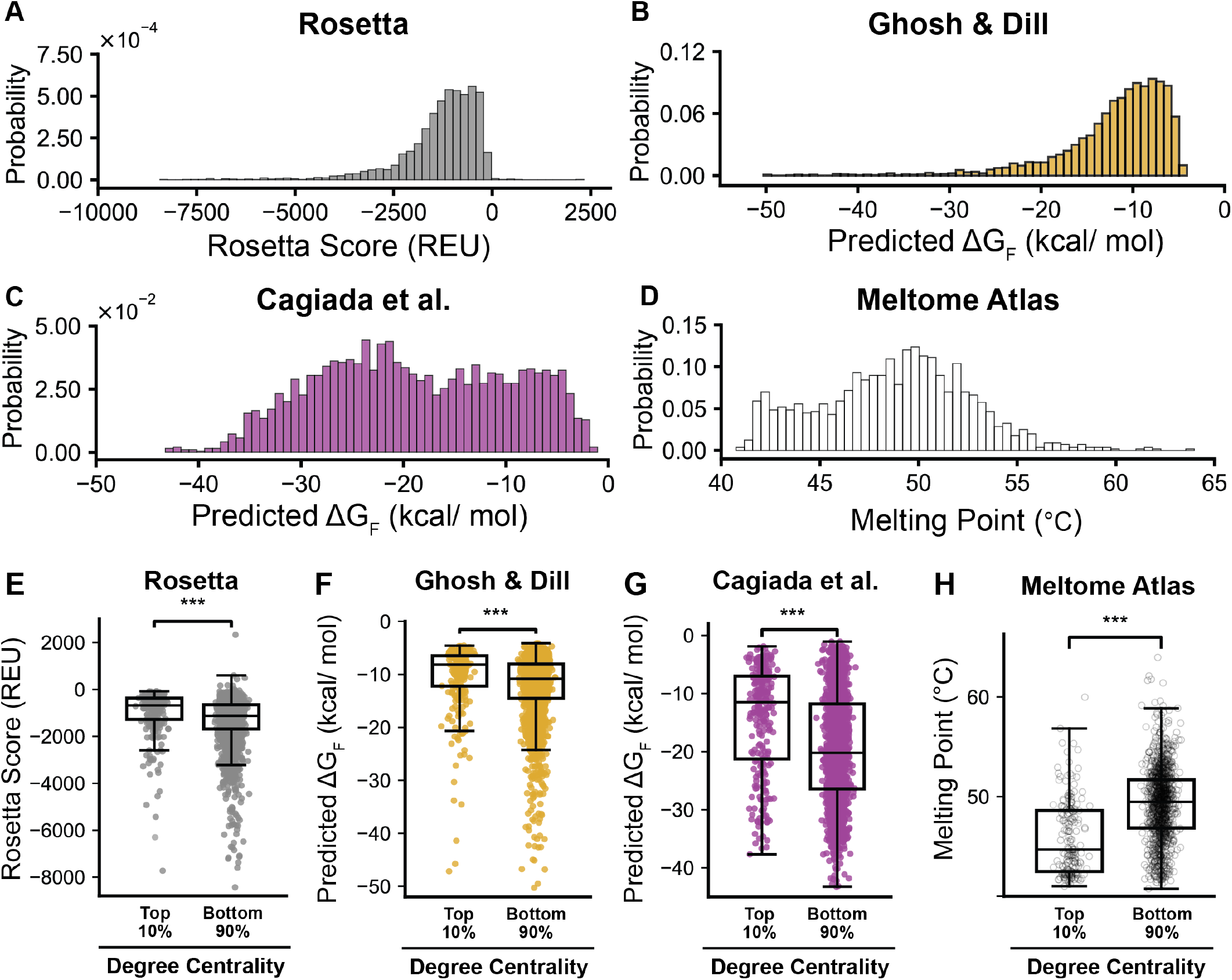
Conformational Stability of Hub Proteins. Experimental and computational estimates of protein stability are compared for hub proteins (≥ 90^th^ percentile degree centrality) and non-hub proteins (< 90^th^ percentile degree centrality). Histograms depict the distribution of free energy of folding (Δ*G*) values for the 2529 water-soluble proteins that adopt a specific fold, according to A) the Rosetta energy function (extreme high-score outliers, B) a length-based approximation, and C) a pre-trained machine-learning tool. D) A histogram depicts the distribution of thermal melting temperatures for the same subset of 2529 water-soluble proteins. E) A box and whisker plot compares the statistical distribution of Rosetta energy scores among hub and non-hub proteins. F) A box and whisker plot compares the statistical distribution of length-based Δ*G* approximations among hub and non-hub proteins. G) A box and whisker plot compares the statistical distribution of machine-learning-based Δ*G* approximations among hub and non-hub proteins. F) A box and whisker plot compares the statistical distribution of thermal melting temperatures among hub and non-hub proteins. The upper and lower edges of the boxes correspond to the 75^th^ and 25^th^ percentile value, respectively. The upper and lower whiskers extend to 1.5 times the interquartile range. Midlines reflect the median value. Statistical significance is indicated by *** (Mann-Whitney U-test, p < 10^−13^).

A comparison of the protein stability distribution across hub proteins (i.e ≥ 90^th^ percentile degree centrality) and the remaining non-hub proteins suggests hub proteins are generally less stable than their interaction partners, regardless of the specific approach used for stability prediction (Mann-Whitney U-test, p < 10^−13^, Fig. 3 E-G). To validate this result, we compared a recently described proteomic melting temperature (T_m_) measurements, which were determined by tracking changes in the solubility of each protein within a yeast lysate as a function of temperature. Remarkably, we find that hub proteins exhibit lower melting temperatures relative to non-hub proteins (Fig. 3H, Mann-Whitney U-test, *p* = 3.8 × 10^−27^). By comparison, bottleneck proteins in the AE-MS network exhibit minimal differences with respect to stability predictions or melting temperature (Fig. S2). Moreover, no significant biases are observed among hub or bottleneck proteins within the Y2H network (Fig. S3). The lack of stability bias in the Y2H network could potentially reflect the lack of native variations in the cellular abundance among Y2H interactors. (see *Discussion*). Taken together, our findings suggest proteins that act as hubs tend to be less stable.

### Predictive Biochemical and Biophysical Features of Hub Proteins

Our collective findings suggest hub proteins exhibit bias with respect to their cellular abundance, turnover, and thermodynamic stability. To determine whether these properties are sufficient to predict whether a protein is likely to act as a hub, we utilized the structural and biochemical features described above to train an ensemble of random forest models to classify soluble, folded proteins as hub (≥ 90^th^ percentile degree centrality) or non-hub proteins. A receiver operating characteristic curve for our top performing model demonstrates hub proteins within the AE-MS network model can be classified with considerable accuracy using only a small collection of twelve features (AUROC = 0.898, Fig. 4A). Performance features for other models trained to classify hubs and bottlenecks in both the AE-MS and Y2H networks can be found in Table S1. Notably, these same features are generally unable to classify bottlenecks within this network (Table S1, Fig. 4A). We were also unable to classify the hubs or bottlenecks within the Y2H network (Table S1, Fig. 4A). A *Shapley Additive exPlanations* (SHAP) analysis of the top performing models for AE-MS hub classifier suggests the two most predictive features are their cellular abundance and half-life (Fig. 4B). By comparison, these features have little value for the Y2H hub classifier model (Fig. 4C). The divergent influence of cellular abundance and half-life measurements very likely arises from distinctions in the underlying experimental designs-AE-MS interactions are measured with endogenous protein mixtures while Y2H measurements are carried out with exogenously expressed proteins. Taken together, our results imply that the tendencies for proteins to form numerous interactions is heavily influenced by their thermodynamic stability and cellular abundance.

**Figure 4.**
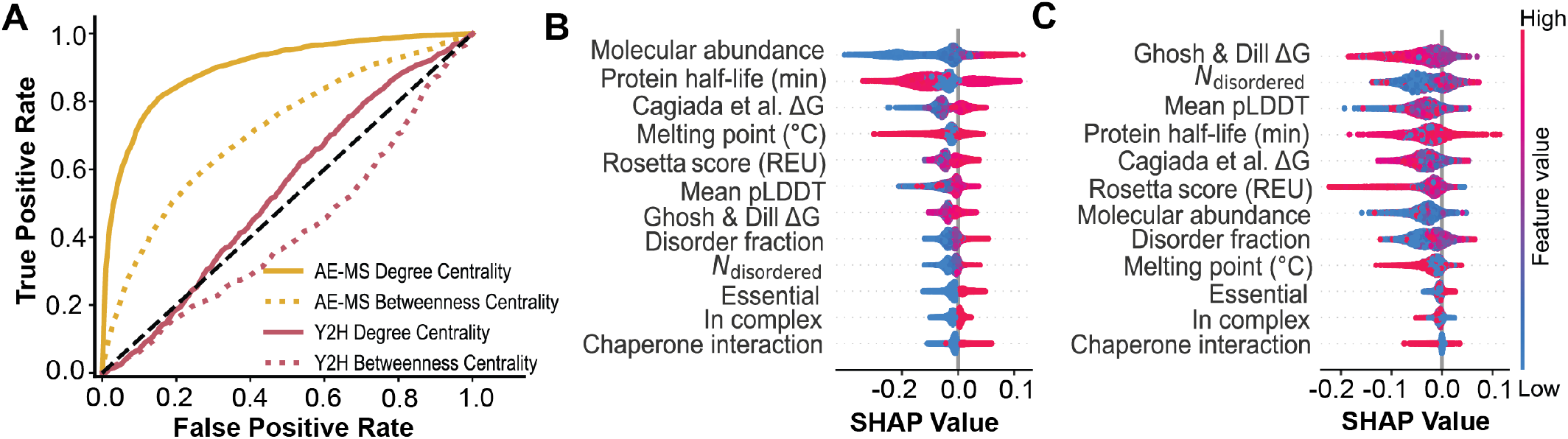
Predictive Features of Network Hubs. Random forest models were trained to classify nodes as hub proteins (≥ 90th percentile degree centrality) and non-hub proteins (< 90th percentile degree centrality) based on various biophysical features and bioanalytical measurements. A) A receiver-operating characteristic diagram plots the true positive rate as a function of the false positive rate for various random forest ensemble models that were trained to classify hub proteins (≥ 90th percentile degree centrality). ROC-AUC values for ensemble random forest models trained to classify hub proteins (≥ 90th percentile degree centrality, solid lines) or bottlenecks (≥ 90th percentile betweenness centrality, dashed lines) in the AE-MS (yellow) or Y2H (pink) network models. B) A summary plot for a SHapley Additives exPlanations (SHAP) analysis of the top-performing model AE-MS hub classifier model depict how individual feature values for each protein factor into the classification output for each node. C) A summary plot for a SHapley Additives exPlanations (SHAP) analysis of the top-performing model Y2H hub classifier model depict how individual feature values for each protein factor into the classification output for each node. The twelve features employed to train each model are arranged from most predictive (top) to least predictive (bottom) based on mean absolute SHAP value. Feature values for each node are colored according to their feature value, where pink values are greater than blue values. On the x-coordinate, negative SHAP values indicate features that weight predictions toward non-hub classification, and vice versa. The horizontal spread and density of dots indicate the distribution and strength of each feature’s effect across the dataset.

### Emergent Properties of Hub Proteins

Proteins that persist within functional complexes along with their binding partners could potentially be occluded from degradative interactions with quality control machinery in a manner that could slow their cellular turnover.^6^ To determine whether variations in steric protection may alter the proteostatic fates of hub proteins, we used the Complex Portal database to parse hub proteins according to whether they reside within stable oligomeric assemblies (static hubs) or whether they instead form a high number of dynamic interactions (dynamic hubs).^37^ Overall, we find that static hubs have a similar median half-life to dynamic hubs despite considerable variability (Fig. 5A, Mann-Whitney U-test, *p* = 0.002).^37^ While trends among stability predictors suggest static hubs are generally more stable than dynamic hubs, the increases in the medians are not statistically significant (Fig. B-D). However, thermal melting measurements clearly show a statistically significant increase in the melting temperature of static hubs relative to dynamic hubs (Fig. 5E, Mann-Whitney U-test, *p* = 7.0 × 10^−4^). Similar trends are not observed among AE-MS bottleneck proteins, nor are they found among Y2H hubs/ bottlenecks (Fig. S4 & S5). In contrast to general trends in hub stability (Fig. 3), these findings suggest that hub proteins that function within stable protein complexes may exhibit higher stability within the cell.

**Figure 5.**
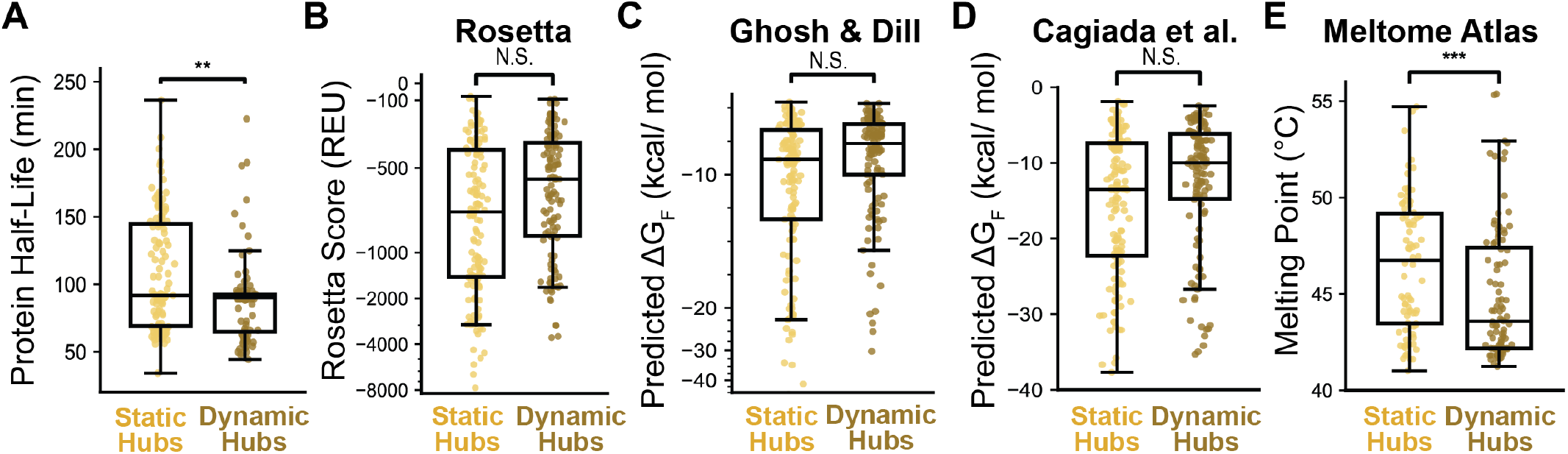
Context-Dependent Properties of AE-MS Hub Proteins. Experimental and computational estimates of protein stability are compared for hub proteins (≥ 90th percentile degree centrality) that function as component of a protein complex (static hubs) or free in solution (dynamic hubs) in the AE-MS network. A) A box and whisker plot compares the statistical distribution of protein half-life measurements among static and dynamic hub proteins. B) A box and whisker plot compares the statistical distribution of Rosetta energy scores among static and dynamic hub proteins. C) A box and whisker plot compares the statistical distribution of length-based ΔG approximations among static and dynamic hub proteins. D) A box and whisker plot compares the statistical distribution of machine-learning-based ΔG approximations among static and dynamic hub proteins. E) A box and whisker plot compares the statistical distribution of thermal melting temperatures among static and dynamic hub proteins. The upper and lower edges of the boxes correspond to the 75^th^ and 25^th^ percentile value, respectively. The upper and lower whiskers correspond to the 90^th^ and 10^th^ percentile value, respectively. The upper and lower whiskers extend to 1.5 times the interquartile range. Midlines reflect the median value. Statistical significance is indicated by ** p < 0.002, *** p < 10^−3^ (Mann-Whitney U-test).

### Chaperone-Hub Interactions are Associated with Kinetic Stability

To explore the mechanistic basis for the divergent turnover rates observed among hub proteins within the yeast interactome (Fig. 2B), we compared the properties of hub proteins that avidly interact with molecular chaperones. We first calculated the fraction of the edges that correspond to chaperone interactions for each node. Overall, we find that the edges associated with hub proteins are statistically enriched with chaperone interactions relative to non-hubs (Fisher’s Exact Test odds ratio = 3.18, *p* = 1.2 × 10^−13^, Fig. 6A), which perhaps stems from their high abundance and low stability. Moreover, dynamic hubs exhibit a significantly higher proportion of chaperone interactions (47.5%) relative to static hubs (19.4%). Indeed, the odds that a dynamic hub interacts with a chaperone are significantly higher than the odds a static hub interacts with a chaperone (Fig. 6B, Fisher’s Exact Test odds ratio = 3.8, *p* = 2.4 × 10^−6^). As expected, the propensity of hubs to interact with chaperones appears to be associated with conformational stability. Trends among stability predictors suggest chaperone-dependent hubs are generally less stable than chaperone-independent hubs, though the increases in the medians are not statistically significant (Fig. 6 D-F). However, chaperone-independent hubs do have significantly higher melting temperatures (Mann-Whitney U-test, *p* = 7.0 × 10^−4^, Fig. 6G). Hub subtypes also exhibit variations in kinetic stability. For instance, though chaperone interactions do not impact static hub turnover, dynamic hubs that exhibit chaperone-dependence have a longer half-life (median value 91.2 min) relative to chaperone-independent dynamic hubs (median value 72.6 min, Mann-Whitney U-test, *p* = 0.001; Fig. 6C). These observations generally suggest that chaperone interactions help extend the lives of unstable hub proteins that form an array of competing interactions.

**Figure 6.**
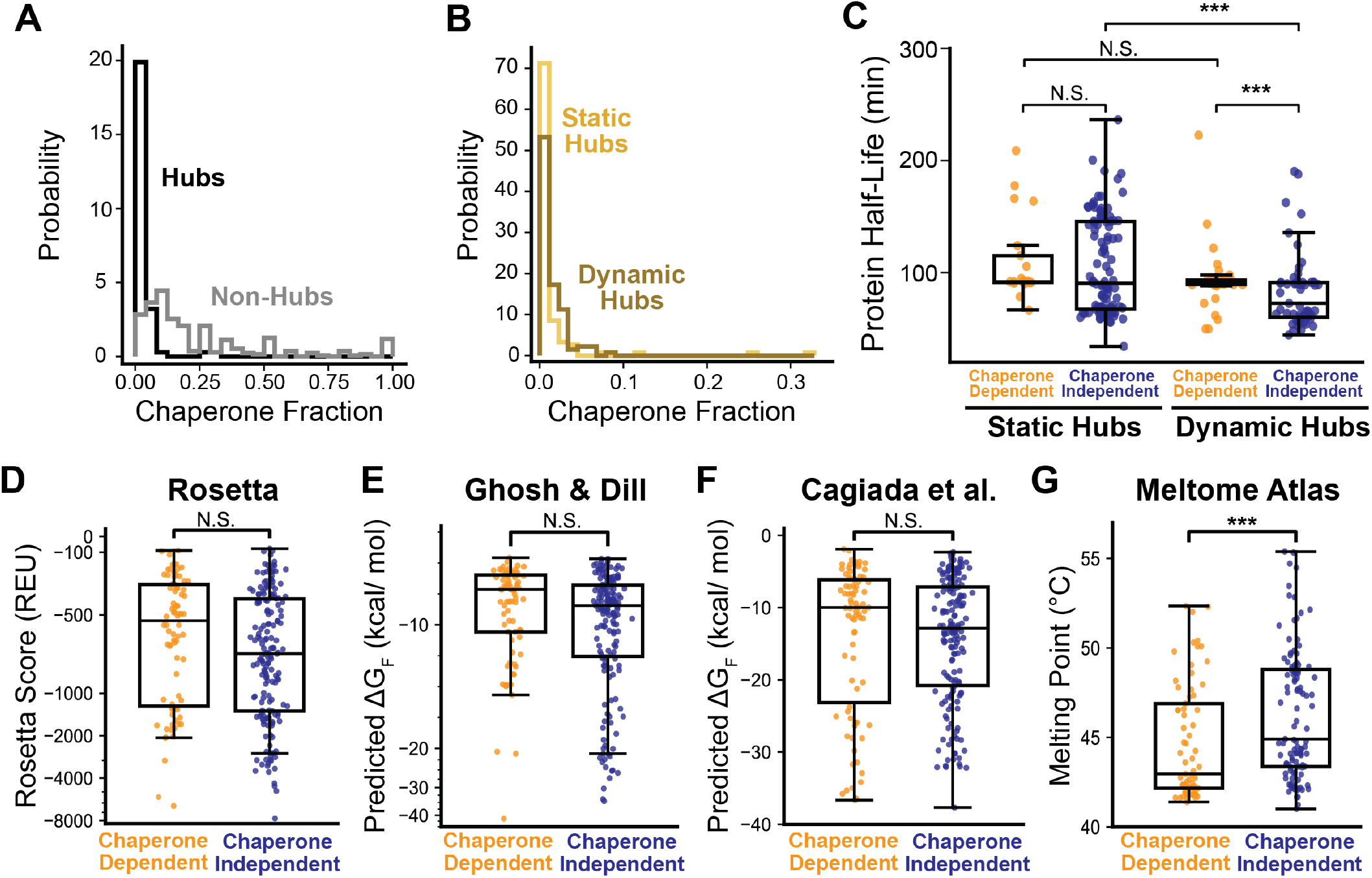
Chaperone Dependence of Hub Proteins. The fraction of edges that constitute interactions with molecular chaperones (chaperone fraction) is analyzed in relation to the stability and turnover of hub proteins in the AE-MS network. A) A histogram compares the fraction of chaperone interactions formed by hub proteins (≥ 90th percentile degree centrality, yellow) and non-hub proteins (< 90th percentile degree centrality, grey). Mann–Whitney U test, *p* = 5.6 × 10^−8^. B) A histogram compares the fraction of chaperone interactions formed by hubs that function as component of a protein complex (static hubs, yellow) or free in solution (dynamic hubs, brown). Mann–Whitney U test, *p* = 3.8 × 10^−5^. C) A box and whisker plot compares the statistical distribution of protein half-life measurements of chaperone-dependent and chaperone-independent hub proteins. D) A box and whisker plot compares the statistical distribution of protein half-life measurements of chaperone-dependent and chaperone-independent static (left) and dynamic (right) hub proteins. E) A box and whisker plot compares the statistical distribution of Rosetta energy scores among chaperone-dependent and chaperone-independent hub proteins. F) A box and whisker plot compares the statistical distribution of length-based ΔG approximations among chaperone-dependent and chaperone-independent hub proteins. G) A box and whisker plot compares the statistical distribution of machine-learning-based ΔG approximations among chaperone-dependent and chaperone-independent hub proteins. H) A box and whisker plot compares the statistical distribution of thermal melting temperatures among chaperone-dependent and chaperone-independent hub proteins. The upper and lower edges of the boxes correspond to the 75^th^ and 25^th^ percentile value, respectively. The upper and lower whiskers extend to 1.5 times the interquartile range. Midlines reflect the median value. Statistical significance is indicated by *** (Mann–Whitney U-test, *p* < 10^−3^).

## Discussion

Like many other biological networks, interactions within the proteome exhibit scaling properties that render them robust to a variety of perturbations. Although understanding the collective behavior of PPI networks is of central importance, many efforts to interpret large-scale interaction datasets rely predominantly on empirical correlations, with limited integration of the physical principles that govern oligomerization. Nevertheless, the recent development of scalable tools to predict the structural and biophysical properties of proteins are providing new opportunities to explore how their collective energetics give rise to their observable scaling. In this work, we merge interactome data with an array of bioanalytical measurements and biophysical approximations to identify combinations of factors that bias the *S. cerevisiae* PPI network. Our findings reveal that network hubs are typically less-stable proteins that are highly abundant, yet quickly turned over in the cell (Figs. 2 & 3). Importantly, we show that these features can be used to accurately predict which proteins serve as network hubs (Fig. 4). We also show that there are at least two classes of hubs that exhibit distinct properties depending on whether they act as a component of a functional protein complex (static hub) or instead form an array of competing interactions (dynamic hubs, Fig. 5). We note that we based these classifications on knowledge of physical complexes that exist within cells, which is distinct from previously disputed^14,42^ classifications based on temporal expression patterns (i.e. “party hubs” vs. “date hubs”).^43^ Finally, we find that the interaction of dynamic hubs with chaperones seems to slow their cellular turnover (Fig. 6). Together, these findings reveal how variations in protein stability shape protein interaction networks.

Many efforts to rationalize PPIs have focused on predicting and interpreting the structural complementarity between proteins within oligomeric complexes. While structural interfaces surely encode specificity,^2,5,44^ we find that this information is not required to infer whether or not a protein is likely to be highly connected within the cellular PPI network. Instead, conformational stability and abundance appear to impose a more general constraint on oligomerization energetics in the cell. Indeed, the simple principles of mass action dictate that abundant proteins should form more oligomers, even if the binding affinity associated with their complexes is relatively weak.^2^ Moreover, unstable proteins that exhibit a lower thermodynamic preference to adopt a single conformation may have access to a broader ensemble of structures that promote interactions.^1,38,39^ Cellular turnover kinetics also appear to provide contextual clues related to the number of interactions that can shield hub proteins from degradation (Fig. 2C). Thus, the behavior of hubs could be viewed as a phenotype that emerges from their abundance, folding energetics, interaction modalities, and proteostatic context rather than from some intrinsic structural feature. Unfortunately, proteomic interactome measurements have not been collected across species, which restricts our ability to assess the generality of our mechanistic findings that emerge from the analysis of the AE-MS network. Nevertheless, the predictive models trained herein make use of a relatively modest set of features that are far more accessible across species. Changes in protein abundance and/ or stability could therefore be used to infer changes in the organization of hubs across divergent PPI networks. Additional investigations are needed to survey how conformational stability is partitioned within divergent interaction networks.

Nearly all of the robust statistical signals identified in our collective analysis of twelve biochemical and biophysical descriptors of the nodes within the yeast interactome emerge in the context of a network constructed from an unusually comprehensive proteomic analysis of the yeast interactome.^15^ By comparison, we observe very few signals within an analogous collection of Y2H measurements. The nature of our findings provides a clear explanation for the divergence of these orthogonal networks. Specifically, our observations concerning the elevated abundance of hubs and bottlenecks highlight the importance of endogenous protein stoichiometries, which are better approximated in the context of AE-MS experiments. Maintaining near-native abundances through the use of cell lysates may preserve protein complexes and recapitulate the certain weak, sub-stoichiometric interactions that cannot be detected with other assays.^21,45^ This perspective helps reconcile inconsistencies in the centrality-lethality rule,^9,13,46^ which posits that essential proteins are highly connected within PPI networks. Others have noted that this relationship only emerges from analyses of certain types of interactome measurements.^47,48^ Our results suggest these discrepancies potentially arise from the divergent properties of static hubs, dynamic hubs, and/ or chaperone-dependent hubs. AE-MS measurements may reveal a wider array of these hubs because they can detect higher order (i.e. non-binary) interactions that form within native complexes composed of static hubs. Nevertheless, while the systematic enrichment of abundant proteins within large-scale AE-MS interactome measurements may catch more of true-positive interactions, the practical consequences of abundance-driven co-purification may also give rise to some number of false-positives.^49^ It is important to bear in mind the caveats of these techniques when interpreting the origins of specific interactions.

Our findings have a variety of implications for the organization and evolution of PPI networks. Hub proteins must find ways to remain highly abundant without gaining the excessive stability needed to escape degradation by QC. One can imagine many ways this could be achieved. Our data suggest many hubs that are not sequestered within static protein complexes may instead become reliant on the protection of molecular chaperones, which are generally also function as hubs. In addition to stoichiometric constraints on biosynthesis,^50^ the fitness landscapes of hub proteins must therefore be constrained by a variety of conformational and proteostatic factors. By comparison, high-betweenness nodes that act as bottlenecks face relatively few of these constraints with the exception of high cellular abundance (Fig. 2B). As is true for hubs, these high-betweenness proteins can also regulate important cellular processes.^13,51-53^ The relative lack constraints on these proteins suggests adaptive changes to network topology may be more readily accomplished through modifications to bottlenecks than to hubs. Additional research is needed to explore how variations in protein stability may drive changes in network architecture.

## Materials and Methods

### Construction of Network Models

Network models were derived from the Cytoscape^54^ session file for The Yeast Interactome, downloaded from www.yeast-interactome.org.^15^ The session file was opened in Cytoscape v3.10.3, and node- and edge-level tables exported via the Cytoscape graphical user interface for downstream use. Weighted k-shell decomposition values were computed for each node using the wk-shell-decomposition app in Cytoscape.^55^ The exported node and edge tables, together with the wk-shell values, served as the primary inputs to the annotation pipeline described below.

Network models were constructed and centrality calculations carried out with NetworkX v3.4.2 in Python.^56^ Degree centrality, betweenness centrality, eigenvector centrality, closeness centrality, load centrality, and PageRank (damping coefficient 0.85) were calculated using the full network of 3,927 nodes, while information centrality was calculated using the largest fully connected subgraph of 3,839 nodes. Yeast two-hybrid network models and centrality calculations were performed identically using data obtained from Center for Cancer Systems Biology Yeast Interactome Database.^13^

### Annotation of Interaction Networks

Protein features were integrated from federated databases (e.g., UniProt, InterPro, Complex Portal) as well as selected high-quality studies. Protein sequences were retrieved from the Saccharomyces Genome Database (SGD)^57^ and mapped to network nodes. Proteins were first classified as globular or transmembrane with DeepTMHMM,^40^ and signal peptide cleavage sites were predicted with SignalP 6.0.^41^ For proteins with predicted signal peptides, the signal peptide region was removed, and all subsequent analyses were performed on these trimmed sequences. Intrinsically disordered regions were predicted using metapredict v3.0^58^ for all proteins for which a sequence was obtained.

For all proteins classified as non-transmembrane by DeepTMHMM, folding free energies were estimated using two approaches: (1) Equation 1 from Ghosh & Dill (2010),^59^ which depends only on protein length and temperature, and (2) the ESM-IF-based model from Cagiada and coworkers^31^ applied to AlphaFold2-predicted structures.^28^ AlphaFold2 predictions (version 4) for the yeast proteome were obtained from the European Bioinformatics Institute (EBI).^60^ For proteins with predicted signal peptides, or where the EBI structure sequence did not match the SGD reference sequence, new AlphaFold2 predictions were carried out. All structures were then relaxed in Rosetta^61^ using the FastRelax protocol with the ref2015 scoring function and ten independent runs per protein; values reported in Fig. 3 correspond to the lowest scoring model across all ten runs.

Finally, additional node-level annotations were integrated from curated sources including protein half-lives,^34,35^ protein abundances,^33^ chaperone interactions,^62^ oligomerization state from Complex Portal,^63^ and essentiality information from the SGD.^64^ A graphical browser that enables exploration of the ANnotated Yeast Interaction network (ANYI) is available on DockerHub.^65^

### Training of Machine Learning Models

The node-level descriptors described above were utilized as features to train an ensemble of random forest models to classify nodes based on their degree centrality or betweenness centrality in the context of the AE-MS or Y2H network. For each network, proteins were categorized as hubs versus non-hubs or bottlenecks versus non-bottlenecks based on whether they are above or below the 10^th^ percentile value. Proteins were then randomly split into training and test sets (80:20). We then used these sets to train and score a series of random forest classifier models with Python using the scikit-learn library. The performance of the resulting models was evaluated using various metrics including balanced accuracy, precision, recall, F1 score, and the area under the receiver operating characteristic curve (AUROC). To interpret the behavior of these models, we used the SHAP library in Python to compute SHapley Additive exPlanation (SHAP) values, which leverage the principles of game theory to quantify the contribution of each feature to individual predictions.

### Statistical Analyses

All statistics were computed in Python using the SciPy (v1.17) library.^66^ Mann-Whitney U tests (scipy.stats.mannwhitneyu) were used for pairwise comparisons shown in Figs. 2, 3, 5, and 6 unless otherwise noted. Bootstrap 95% confidence intervals were computed using custom Python code with 10^6^ resamples in all cases. Complete, executable examples of the bootstrapping procedure are available in the GitHub repository. *p*-values were adjusted using the Benjamini–Hochberg procedure within four pre-specified families defined by centrality measure (degree or betweenness) and hypothesis type (Mann–Whitney U tests for quantitative comparisons; Fisher’s Exact Tests for categorical enrichments).

## Data Availability

All raw data used in this study are publicly available on the CyVerse Data Store at [/iplant/home/shared/NCEMS/working-groups/energetic-origins/required-data]. A free CyVerse account is required to download the data. All Python code used for network annotation and analysis is available on GitHub at [https://github.com/NCEMS/energetic-origins-of-PPI-connectivity.git], together with the processed dataset and the Jupyter notebooks used to generate all main-text and supplementary figures.

## Acknowledgements

We thank Eugene I. Shakhnovich for scientific input. This work was supported and coordinated by the National Science Foundation (NSF) through the National Synthesis Center for Emergence in the Molecular and Cellular Sciences (NCEMS) under Grant NSF MCB-2335029. A.W.R.S. acknowledges support from the Canadian Institute of Health Research grant MOP-G-408523, Natural Sciences and Engineering Research Council of Canada grant RGPIN-2016-06566, and the Canada Research Chairs. X.C.G. acknowledges support from the Natural Sciences and Engineering Research Council of Canada – Masters Canada Graduate Scholarship and Fonds de Recherche du Québec en Santé – Masters Training Scholarship. This work was also supported by NSF award IOS-2038872, and OIA-2418230 to M.S.M as well as by the National Institute of General Medical Sciences (R35GM152086) to J.P.S.

