## Supplemental Materials for "Protein Stability, Turnover Kinetics, and Abundance Constrain the Scaling of Protein Interaction Networks"

‡Authors contributed equally

*Corresponding Authors: jschleba (at) purdue.edu, adrian.serohijos (at) umontreal.ca, mshahid (at) clemson.edu

*This File Includes:*
Figure S1
Figure S2
Figure S3
Figure S4
Figure S5
Table S1

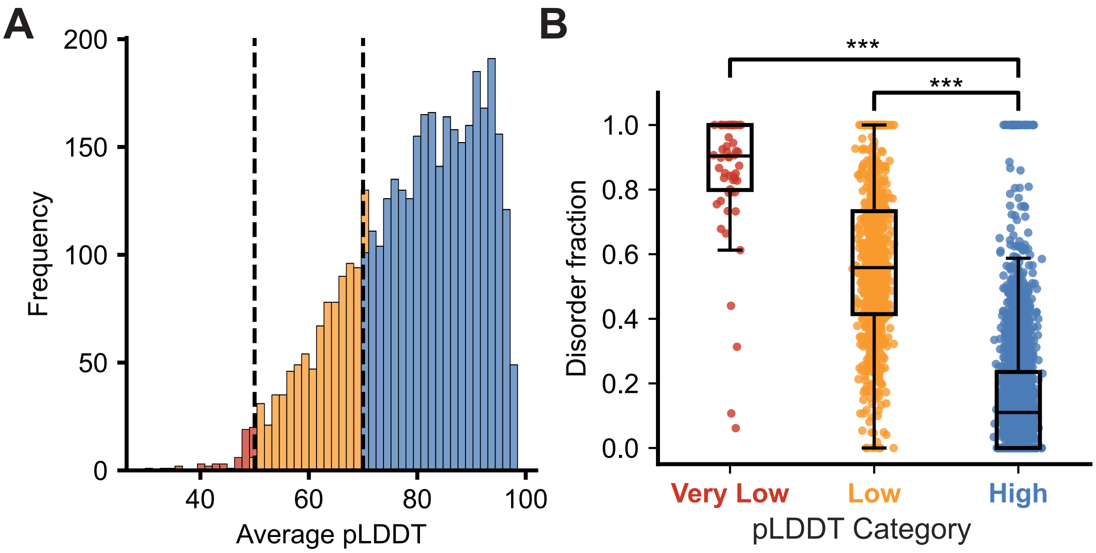


**Figure S1. Distribution of pLDDT scores in AlphaFold2 models and their relation to protein disorder in the yeast proteome.** A) A histogram depicts the distribution of average pLDDT scores across AlphaFold2 models of nodes within the PPI network. Scores were used to classify structural models as either very low-confidence (average pLDDT< 50, red), low-confidence (average pLDDT = 50-69, orange), or high-confidence (average pLDDT ≥ 70, blue). B) Metapredict v3.0 was used to calculate the fraction of disordered residues within each protein. A box and whis-ker plot depicts how the disordered fraction varies among proteins with very low-confidence, low-confidence, and high-confidence AlphaFold2 models.*** (Mann-Whitney U-test, $p < 0.001$).

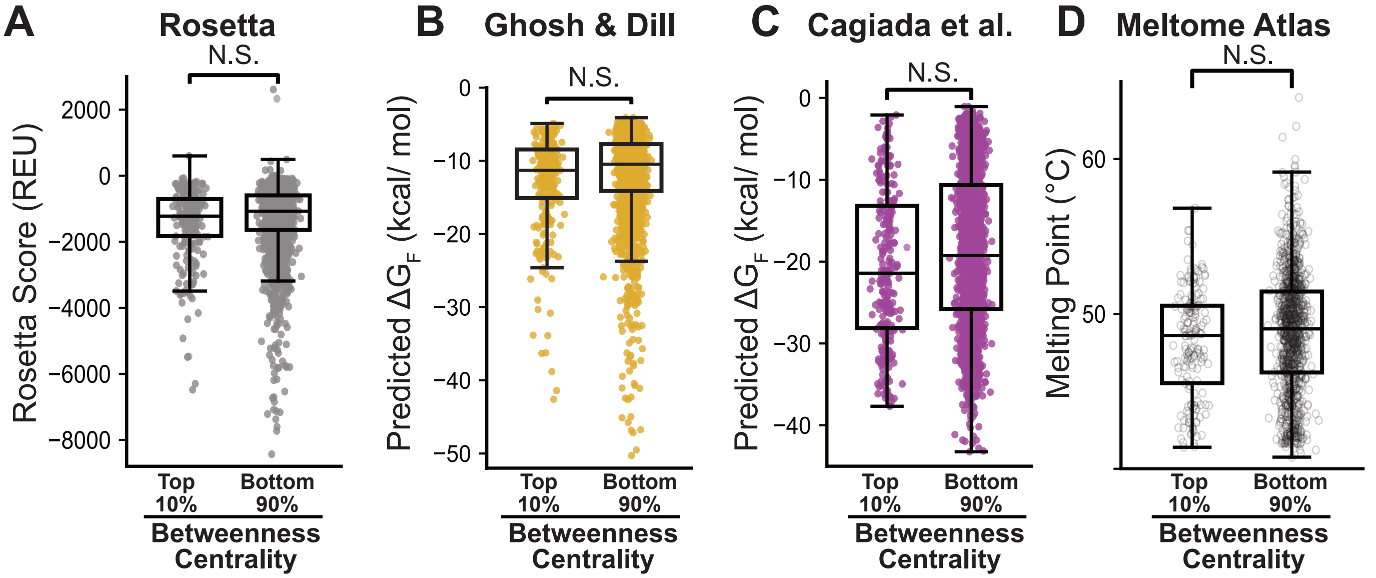


**Figure S2. Conformational stability for bottleneck proteins in the AE-MS network.** A) Box and whisker plots compares the distribution of predicted protein stability values for bottleneck (>90$^{th}$ percentile betweenness centrality) and non-bottleneck (<90$^{th}$ percentile betweenness centrality) proteins in the AE-MS network according to A) the Rosetta energy function (grey), B) length-based approximations (gold), and C) a pre-trained machine learning tool (purple). D) A box and whisker plot compares the thermal melting temperatures (white) bottleneck (>90$^{th}$ percentile betweenness centrality, red) and non-bottleneck (<90$^{th}$ percentile betweenness centrality, blue) proteins in the AE-MS network. Difference were found to be not statistically significance according to a Mann-Whitney U-test ($p$ > 0.05).

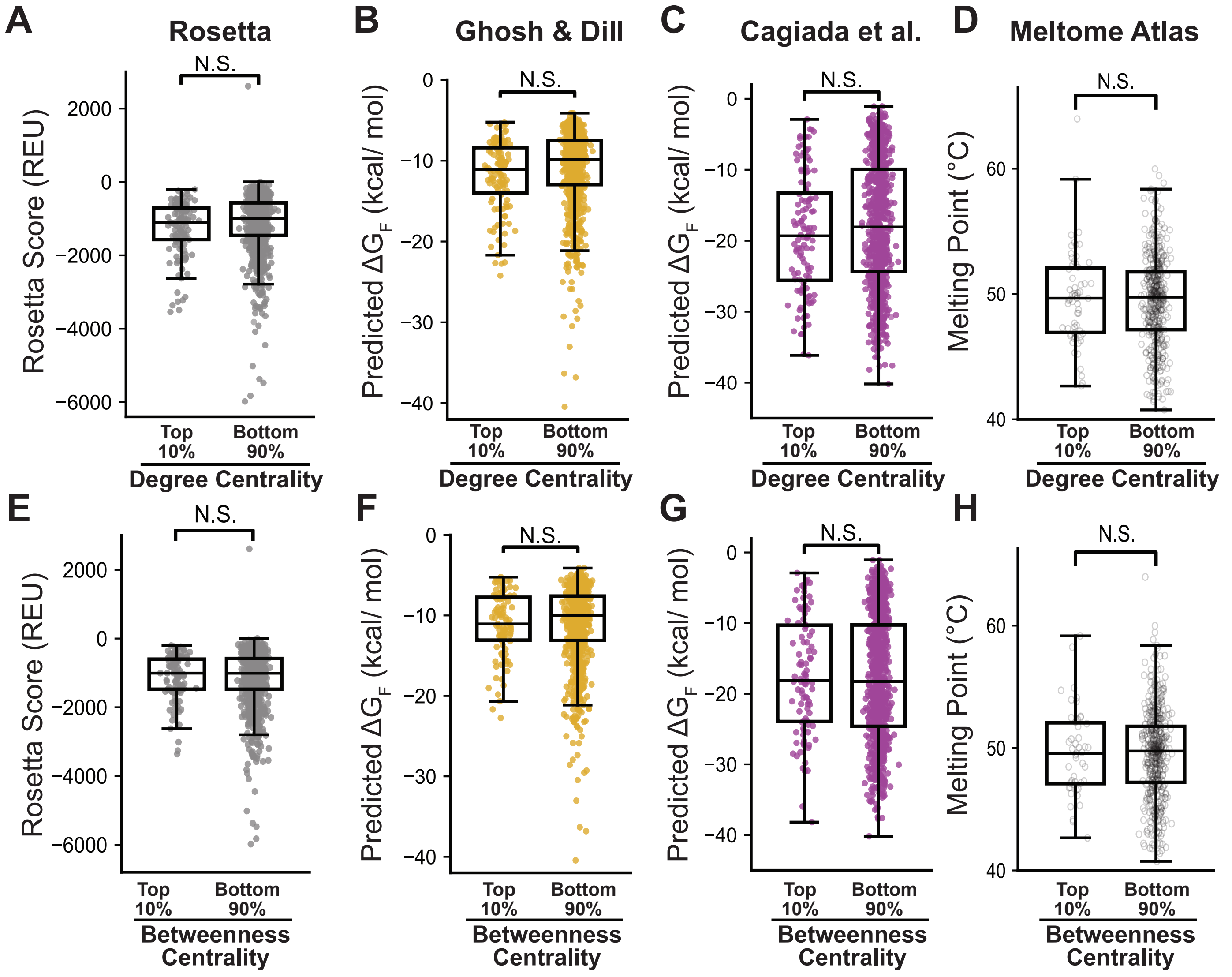


**Figure S3. Conformational stability for hub and bottleneck proteins in the Y2H network.** Box and whisker plots compare the distributions of predicted protein stability values for hub (>90$^{th}$ percentile degree centrality) and non-bottleneck (<90$^{th}$ percentile degree centrality) proteins in the Y2H network according to A) the Rosetta energy function (grey), B) length-based approximations (gold), and C) a pre-trained machine learning tool (purple). D) A box and whisker plot compares the thermal melting temperatures (white) bottleneck (>90$^{th}$ percentile betweenness centrality) and non-bottleneck (<90$^{th}$ percentile betweenness centrality) proteins in the Y2H network. A second set of box and whisker plots compare the distributions of predicted protein stability values for bottleneck (>90$^{th}$ percentile betweenness centrality) and non-bottleneck (<90$^{th}$ percentile betweenness centrality) proteins in the Y2H network according to E) the Rosetta energy function (grey), F) length-based approximations (gold), and G) a pre-trained machine learning tool (purple). H) A box and whisker plot compares the thermal melting temperatures (white) bottleneck (>90$^{th}$ percentile betweenness centrality) and non-bottleneck (<90$^{th}$ percentile betweenness centrality) proteins in the Y2H network. Difference were found to be not statistically significance according to a Mann-Whitney U-test ($p > 0.05$).

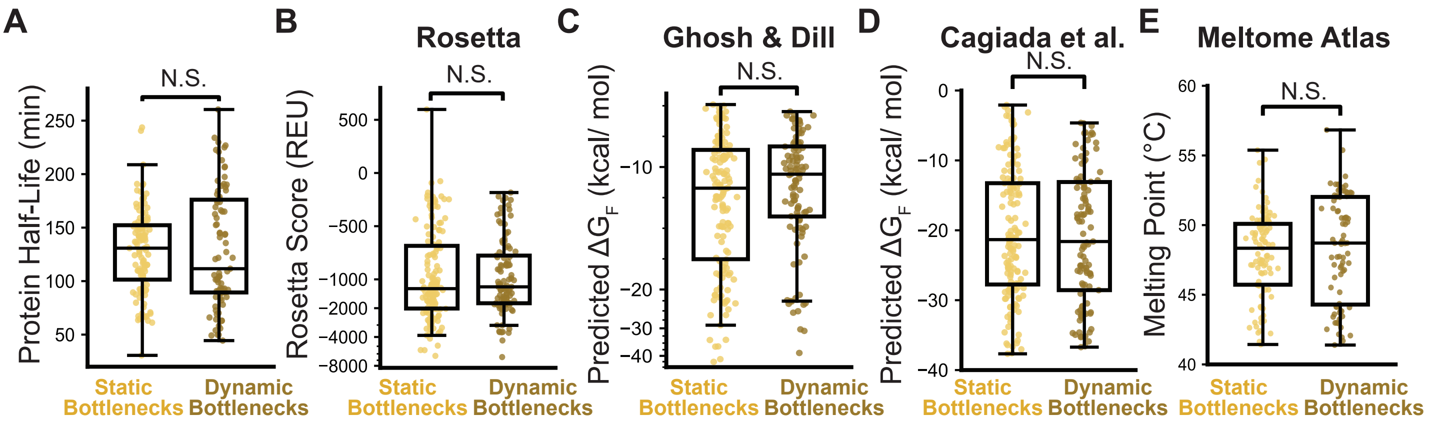


**Figure S4. Context-Dependent Properties of AE-MS Bottleneck Proteins.** Experimental and computational estimates of protein stability are compared for bottleneck proteins (≥ 90th percentile betweenness centrality) that function as component of a protein complex (static hubs) or free in solution (dynamic hubs) in the AE-MS network. A) A box and whisker plot compares the statistical distribution of protein half-life measurements among static and dynamic hub proteins. B) A box and whisker plot compares the statistical distribution of Rosetta energy scores among static and dynamic hub proteins. C) A box and whisker plot compares the statistical distribution of length-based Δ*G* approximations among static and dynamic hub proteins. D) A box and whisker plot compares the statistical distribution of machine-learning-based Δ*G* approximations among static and dynamic hub proteins. E) A box and whisker plot compares the statistical distribution of thermal melting temperatures among static and dynamic hub proteins. The upper and lower edges of the boxes correspond to the 75th and 25th percentile value, respectively. The upper and lower whiskers extend to 1.5 times the interquartile range. Midlines reflect the median value. Difference were found to be not statistically significance according to a Mann-Whitney U-test ($p > 0.05$).

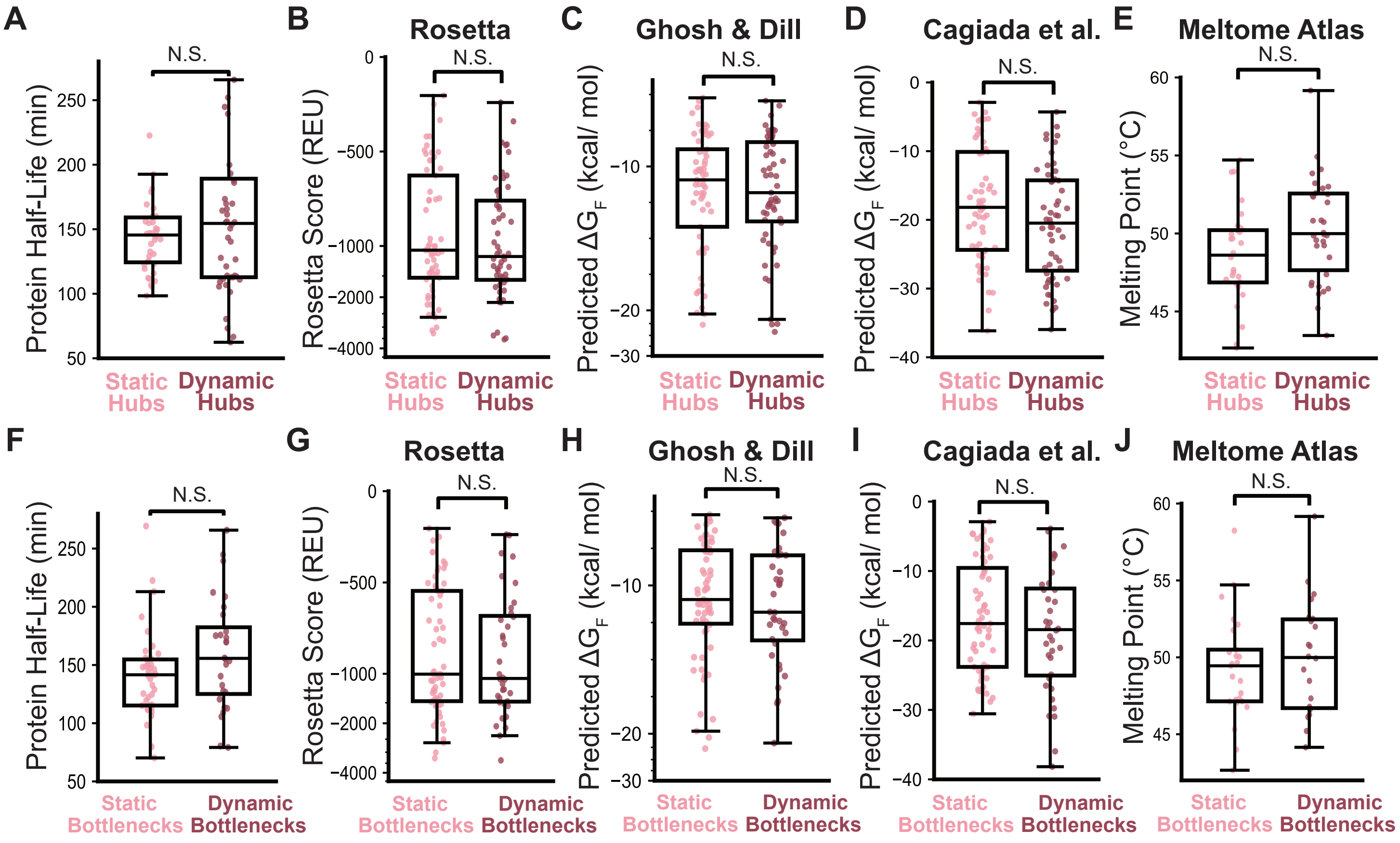


**Figure S5. Context-Dependent Properties of Y2H Hubs and Bottleneck Proteins.** Experimental and computational estimates of protein stability and turnover are compared among hub proteins (≥ 90$^{th}$ percentile degree centrality) and bottleneck proteins (≥ 90$^{th}$ percentile betweenness centrality) that function as component of a protein complex (static hubs) or free in solution (dynamic hubs) in the Y2H network. Box and whisker plots compare the statistical distribution of protein half-life measurements among static and dynamic A) hub proteins or F) bottleneck proteins. Box and whisker plots compare the statistical distribution of Rosetta energy scores among static and dynamic B) hub proteins or G) bottleneck proteins. Box and whisker plots compare the statistical distribution of length-based $\Delta G$ approximations among static and dynamic C) hub proteins and H) bottleneck proteins. Box and whisker plots compare the statistical distribution of machine-learning-based $\Delta G$ approximations among static and dynamic D) hub proteins and I) bottleneck proteins. Box and whisker plots compare the statistical distribution of thermal melting temperatures among static and dynamic F) hub proteins and J) bottleneck proteins. The upper and lower edges of the boxes correspond to the 75$^{th}$ and 25$^{th}$ percentile value, respectively. The upper and lower whiskers extend to 1.5 times the interquartile range. Midlines reflect the median value. Difference were found to be not statistically significance according to a Mann-Whitney U-test ($p > 0.05$).

### Table S1. Supervised Model Performance

*Models trained to classify hubs and bottlenecks in both the AE-MS and Y2H networks w/ the feature set (n= 12).*

| Model | Balanced Accuracy | F1-Score | Precision | Recall | ROC-AUC |
|---|---|---|---|---|---|
| **AE-MS degree centrality** | 0.717 | 0.545 | 0.675 | 0.460 | 0.898 |
| **AE-MS betweenness centrality** | 0.560 | 0.207 | 0.342 | 0.151 | 0.713 |
| **Y2H degree centrality** | 0.496 | 0.004 | 0.017 | 0.003 | 0.536 |
| **Y2H betweenness centrality** | 0.500 | 0.000 | 0.000 | 0.000 | 0.421 |
